# ACTB and GAPDH proteins appear at multiple positions of SDS-PAGE and may not be suitable for serving as reference genes for the protein level determination in such techniques as Western blotting

**DOI:** 10.1101/2020.03.05.978494

**Authors:** Yan He, Ju Zhang, Jiayuan Qu, Lucas Zellmer, Yan Zhao, Siqi Liu, Hai Huang, Dezhong Joshua Liao

## Abstract

Most human genes can produce multiple protein isoforms that should appear at multiple positions of polyacrylamide gel electrophoresis (PAGE) with sodium dodecyl sulfate (SDS), but most published results of Western blotting show only one protein. We performed SDS-PAGE of proteins from several human cell lines, isolated the proteins at the 72-, 55-, 48-, 40-, and 26-kD positions, and used liquid chromatography and tandem mass spectrometry (LC-MS/MS) to determine the protein identities. Although ACTB and GAPDH are 41.7-kD and 36-kD proteins, respectively, LC-MS/MS identified peptides of ACTB and GAPDH at all of these SDS-PAGE positions, making us wonder whether they produce some unknown protein isoforms. The NCBI (National Center for Biotechnology Information, USA) database lists only one ACTB mRNA but five GAPDH mRNAs and one non-coding RNA. The five GAPDH mRNAs encode three protein isoforms, while our bioinformatic analysis identified a 17.6-kD isoform encoded by the non-coding RNA. All LC-MS/MS-identified GAPDH peptides at all positions studied are unique, but some of the identified ACTB peptides are shared by ACTC1, ACTBL2, POTEF, POTEE, POTEI, and POTEJ. ACTC1 and ACTBL2 belong to the ACT family with great similarities to ACTB in protein sequence, whereas the four POTEs are ACTB-containing chimeric genes with the C-terminus of their proteins highly similar to ACTB. These data collectively disqualify GAPDH and ACTB from serving as the reference genes for determination of the protein level in such techniques as Western blotting, a leading role these two genes have been playing for decades in the biomedical research.

## Introduction

In 2012, we reported a bioinformatic study showing that ACTB and GAPDH genes in the human and mouse genomes have large numbers of intronless pseudogenes that are highly similar to the mRNA sequence of ACTB or GAPDH [1]. Since it is a general belief that the whole genomes are basically transcribed to RNA [2], transcripts of these pseudogenes, which are located on different chromosomes and thus are likely swayed by different physiological or pathological conditions, may be mistakenly detected along with the authentic ACTB or GAPDH RNA in RT-PCR (reverse transcription followed by polymerase chain reactions). We therefore suggested that biomedical researchers should take extra caution when using these two genes as references in RT-PCR. Besides this pseudogene issue, ACTB and GAPDH have other weaknesses as the reference genes for normalizing the loading of RNA or protein samples, which has been frequently discussed in the literature [3–8]. Chief among these weaknesses of the GAPDH are its versatile functions, including membrane fusion, apoptosis, regulation of stability and transcription of RNA, instability and repair of DNA, etc., besides its canonical role in energy production [3–5]. Moreover, ACTB has been reported to form fusion genes in some human neoplasms [9–13], while fusion genes involving GAPDH has been reported in evolutionarily low organisms [14–18].

It is well known that most genes in the human genome and in the genomes of many evolutionarily complex species of animals are expressed to multiple protein isoforms to meet various needs in different developmental, physiological, or pathological situations. Actually, each gene is no longer considered to encode a specific phenotype but is perceived to encode a full range of phenotypes. This newly emerging concept, dubbed by Heng et al as “fuzzy inheritance” [19– 21], may be mechanistically attributed in part to the protein multiplicity of each gene. The mechanisms for protein multiplicity are multiple, including alternative transcriptional initiation or termination to produce different RNA transcripts with longer or shorter 5’- or 3’-ends, alternative splicing of a transcript to produce different mRNA variants with more or fewer exons, and alternative uses of translational start or stop codons in a given mRNA to produce different protein isoforms with a longer or shorter N- or C-terminus. Moreover, different individuals may have different single nucleotide polymorphisms, which may affect transcription, splicing, or translation. Many different genetic alterations such as single nucleotide mutations can also affect protein multiplicity in pathological situations by forming fusion genes or by affecting the aforementioned mechanisms.

There currently is still lacking a simple but high-throughput technical approach to determine protein isoforms. High-throughput determination of protein expression is often achieved using a bottom-up procedure of LC-MS/MS (liquid chromatography together with tandem mass spectrometry), in which proteins are first enzymatically digested to short peptides before a LC-MS/MS procedure. The resulting MS data of each short peptide is then matched to a database of protein reference, which results in the identity, i.e. the amino acid (AA) sequence, of the peptide, and in turn the gene’s protein product to which this peptide belongs. Because this procedure uses a short peptide to predict the existence of a whole protein, it is referred to as “bottom-up”. Several years ago, we developed an oversimplified top-down LC-MS/MS strategy, in which proteins were first stratified based on their molecular weights using polyacrylamide gel electrophoresis (PAGE) in the presence of sodium dodecyl sulfate (SDS), followed by isolation of the proteins from the gel at a given position of SDS-PAGE. These proteins with known molecular weights in the SDS-PAGE gel were then subjected to a routine LC-MS/MS procedure for their identification [22]. Our studies with this oversimplified top-down approach surprisingly show that most proteins identified are not supposed to appear at a given position of SDS-PAGE because their theoretical molecular masses are either much too large or much too small. Since Western blotting (WB) in most, if not all, cases can detect proteins at the expected position of SDS-PAGE, we concluded that most of those proteins detected unexpectedly at a given SDS-PAGE position might be additional isoforms besides the canonical or the wild type (WT) form [22–24]. For instance, ACTB and GAPDH are proteins of about 41.7 kD and 36 kD, respectively, and they can indeed be detected at the corresponding position of SDS-PAGE using WB in probably all published studies. However, we detected peptides of ACTB and GAPDH roughly at the 72-kD, 55-kD, 48-kD, 40-kD, and 26-kD positions of SDS-PAGE. We herein report these data of ACTB and GAPDH and discuss the meaning behind the data.

## Materials and methods

### Protein sample preparation and SDS-PAGE

This study contained two separate experiments of SDS-PAGE and LC-MS/MS, as described previously [22,23]. In the first experiment [22], human breast cancer cell line MDA-MB231 (MB231) and human embryonic kidney cell line HEK293 were cultured as routine at 37 C in an incubator with 5% CO_2_ in 10-cm dishes with a DMEM medium containing 10% bovine fetal serum. In the second experiment, human breast cancer cell lines MB231 and MCF7 were cultured in the same way. Cells at 80% confluence were washed with 1x phosphate buffered saline (PBS) and then scraped in a lysis buffer [25] that contained 1× Protease inhibitor cocktail (Sigma-Aldrich, Inc, St. Louis, MS, USA), as described before [22,26]. After the cell lysate was centrifuged at 12,000 rpm for 20 minutes at 4 C, the supernatant was collected as the protein sample and evaluated for protein concentration with a BCA (bicinchonic acid) kit (Pierce, Rockford, IL, USA).

The protein samples were diluted with a gel-loading buffer routinely used for WB, which contained 2% of SDS and 2% of 2-mercaptoethanol in the final concentration. After boiling for 4 minutes and then rapid cooling on ice, the proteins were loaded into a 10% SDS-containing polyacrylamide gel. To better separate and better detect the proteins, the gel was made with 10×10.5 cm glass plates included in the Hoeffer SE260 vertical slab gel system (Hoeffer Inc; http://www.hoeferinc.com/), which produced a gel 2 cm longer in the vertical direction than all gels made using the regular mini-gel cast systems of Hoeffer and other companies. In the first experiment, the 1^st^ well of the gel was loaded with 100 μg (from HEK293 cells) or 70 μg (from MB231 cells) of proteins, the 2^nd^, 3^rd^, and 10^th^ wells were loaded with a pre-stained protein marker that contained bands at 135-kD, 100-kD, 72-kD, 55-kD, 40-kD, 33-kD, 24-kD, and 11-kD, respectively (Fig 1A). The remaining 4^th^-9^th^ wells were loaded with 60 μg of proteins per well. One gel was used for the protein sample from the MB231 cells while another gel was used for the sample from the HEK293 cells. The two gels were electrophoresed simultaneously using the same power supply, and electrophoresis was stopped when the 11-kD marker ran out of the gel. In the second experiment, the first and last wells of the gel were loaded with a pre-stained protein marker while each remaining well was loaded with 50 μg of protein sample (Fig 1B). One gel was loaded with proteins from the MCF7 cells while another gel was loaded with proteins from the MB231 cells. Electrophoresis of the proteins was performed as described above.

**Fig. 1:**
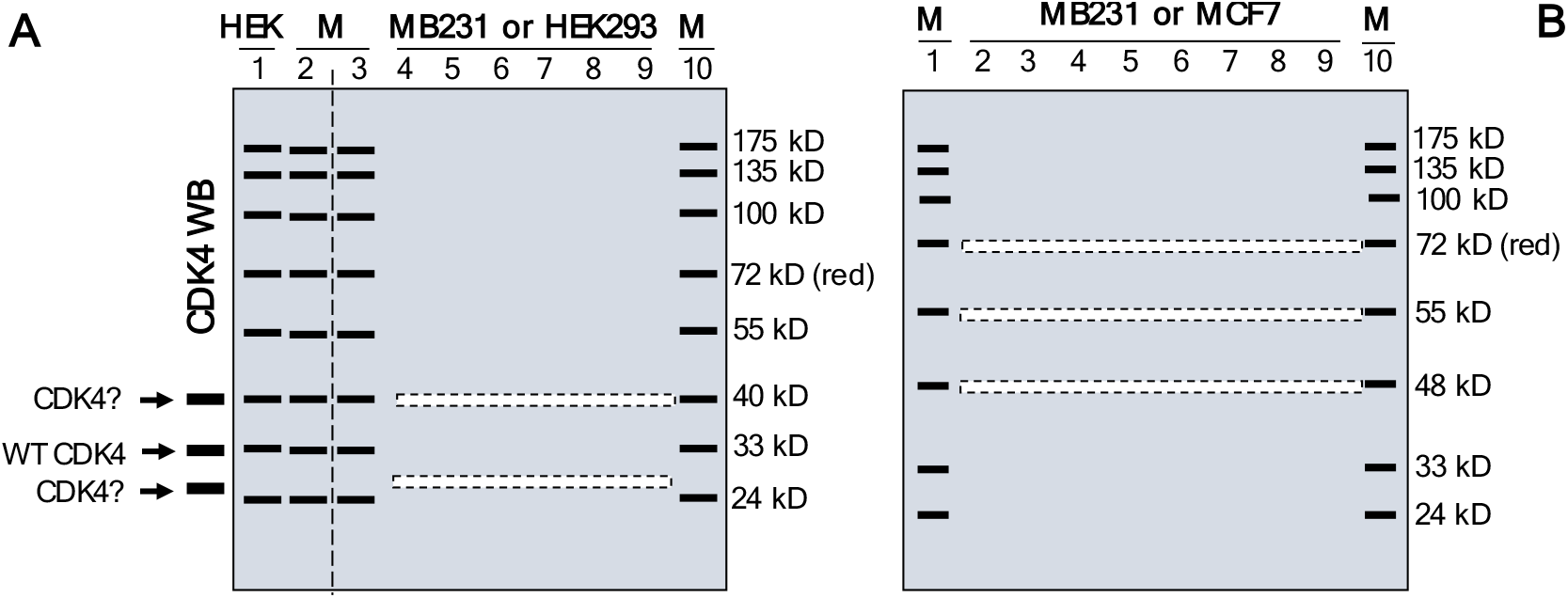
Illustration of excision of narrow stripes of gel after SDS-PAGE, with detail described in the Materials and Methods section. HEK: protein sample from the HEK231 cells. M: pre-stained protein marker.

### Excision of narrow stripes of gel

In the first experiment, the gel was cut vertically with a surgical blade along the dashed line between the 2^nd^ and 3^rd^ lanes as illustrated in Figure 1A. The proteins in the 1^st^ lane and the pre-stained marker in the 2^nd^ lane were then transferred onto a PVDF (polyvinylidene difluoride) membrane via electrophoresis, followed by the detection of CDK4 proteins via a quick WB procedure using the sc-601 primary antibody from Santa Cruz Biotech, as detailed before [27]. The WB was performed in a shorter-than-usual time period and, meanwhile, the remaining part of the gel was stored at 4°C. As we previously reported [27], the WB using several primary antibodies resulted in several bands on the membrane which, besides the WT CDK4 protein exactly at the 33-kD position, included a band at 40-kD and a band slightly higher than the 24-kD band of the pre-stained protein marker, estimated as 26-kD [27]. We then aligned the WB membrane with the remaining part of the gel to determine the 26-kD-band position of the gel. Guided by two rulers along with the pre-stained marker at the 3^rd^ and 10^th^ lanes, we excised out a narrow stripe, about 2 mm in width, at the 26-kD of the 4^th^-9^th^ lanes of each gel, and then another narrow stripe at the 40-kD position shown by the pre-stained marker (Fig. 1A). The narrow gel stripes were illustrated as dashed boxes in Figure 1A, and were used for LC-MS/MS. The initial reason to excise proteins at these positions was to determine whether the WB-detected proteins at these positions were CDK4 isoforms [27].

In the second experiment, a 2-mm stripe of gel was excised at the 72-, 55-, and 48-kD positions, respectively, along the pre-stained protein markers in the first and last lanes. These positions were selected after carefully considering many technical issues: first, we had pre-stained protein markers showing these positions, which allowed us to precisely excise a narrow gel stripe at the correct molecular weight. Second, this 48-72-kD range resides in the middle of the 10% gel cast of most commonly used mini-gel-cast systems. This middle range still leaves us with large regions below 48-kD and above 72-kD. Third, proteins with very large molecular weights, say larger than 150 kD, cannot be well separated in a 10% gel.

### LC-MS/MS

As described before [22,23], the excised gel stripes containing proteins were dehydrated with escalating concentrations of acetonitrile (ACN) as per routine methods. The in-gel proteins were reduced and alkylated with 10 mM dithiothreitol and 55 mM iodoacetamide, followed by digestion with trypsin at 37 °C for 16 hours (5). The tryptic peptides were then extracted from the gel with ACN containing 0.1% formic acid (FA), vacuum-dried, and dissolved in 0.1% FA. The peptides were delivered onto a nano RP column (5-μm Hypersil C18, 75 mm × 100 mm; Thermo Fisher Scientific, Waltham, MA, USA) and eluted with escalating (50-80%) ACN for 60 min at a speed of 400 nL/min. Different fractions of the eluate were injected into a Q-Executive mass spectrometer (Thermo Fisher Scientific, Waltham, MA, USA) set in a positive ion mode and a data-dependent manner with a full MS scan from 350 to 2,000 m/z. High collision energy dissociation (HCD) was used as the MS/MS acquisition method. Raw MS/MS data were converted into an MGF format using Proteome Discoverer 1.2 (Thermo Fisher Scientific, Waltham, MA, USA). The exported MGF files were searched with Mascot v2.3.01 in a local server against the mouse SwissProt database. All searches were performed with a tryptic specificity allowing one missed cleavage. Carbamidomethylation was considered as fixed modification whereas oxidation (M) and Gln->pyro-Glu (N-term Q) were considered variable modifications. The mass tolerance for MS and MS/MS was 15 ppm and 20 mmu, respectively. Proteins with false discovery rates (FDR) < 0.01 were further analyzed.

### Bioinformatic information retrieval and analyses

The information on RNA and protein sequences was retrieved in September of 2019 from the NCBI (National Center for Biotechnology Information, USA) website (https://www.ncbi.nlm.nih.gov/gene/). The open reading frame (ORF) of an RNA and molecular weight of the protein encoded by the ORF were determined using DNAstar software. Sequence alignment was performed using the Blast function of NCBI. Distance tree analysis of RNA sequences was also performed using the Blast function.

### Calculation of total coverage rate and unique coverage rate

The LC-MS/MS procedure generated two basic sets of datasheets, annotated as “proteingroups” and “psms”, respectively. The “proteingroups” datasheet contains “coverage” data (column D in the Supplementary table 1), which is the ratio of the total number of AAs in all LC-MS/MS-identified peptides to the total number of the AAs in the annotated protein of a particular gene. This coverage is coined herein as “the total coverage rate”. The sequence of each identified peptide is given in the “psms” datasheets (Supplementary tables 2 and 3). For many genes, including ACTB, some LC-MS/MS-identified peptides are not unique to the annotated protein of the particular gene but, instead, are also shared by protein(s) of one or more other genes, which are referred to as “common peptides”. We retrieved the sequence of each identified peptide, common or unique, from the “psms” datasheet for GAPDH and ACTB, and mapped the sequence onto the full-length protein of GAPDH or ACTB. We then calculated the total coverage rate, which is the ratio of the total AAs of both common and unique peptides to the total AAs of the full-length GAPDH or ACTB protein. We also calculated the “unique coverage rate”, which is the ratio of the total AAs of the unique peptides to the total AAs in the full-length GAPDH or ACTB protein. A higher unique coverage rate indicates a higher possibility of the presence of the protein in the studied position of the SDS-PAGE gel.

## Results

### RNAs and proteins of GAPDH and ACTB from NCBI

NCBI database lists six RNA variants of the human GAPDH gene, including five normalized mRNA variants annotated as NM_ sequences and one normalized non-coding RNA annotated as a NR_ sequence. Five of the six, including the non-coding one, are derived from alternative splicing of the same transcript, while the remaining one is derived from an alternative initiation of transcription at the first intron of the NM_001289746.2 sequence (Fig 2, top panel). Three proteins (NP_00127665.1, NP_001276674.1, and NP_002037.2 encoded, respectively, by the NM_001289746.2, NM_001289745.3, and NM_002046.7) have the same AA sequence, with the NP_00127665.1 protein shown as a representative in the middle panel of Figure 2, which is considered herein as the full-length protein. Compared with this full-length sequence, protein NP_1344872.1 lacks 18 AAs because its exon 4 is shorter (Fig. 2, top panel), whereas protein NP_001234728.1 lacks the N-terminal 42 AAs because the alternative initiation of transcription leads to alternative usage of translation start codon (Fig. 2, top and middle panels). Although NR_152150.2 is annotated by NCBI as a non-coding RNA, our bioinformatic analysis identified an ORF encoding a GAPDH protein isoform of 161 AAs, which is constituted by the N-terminal 142 AAs and the C-terminal 19 AAs of the full-length protein. Therefore, the human GAPDH gene has at least four protein isoforms based on the NCBI information, with their similarities and disparities as well as their theoretical molecular masses shown in the bottom panel of Figure 2.

**Fig.2:**
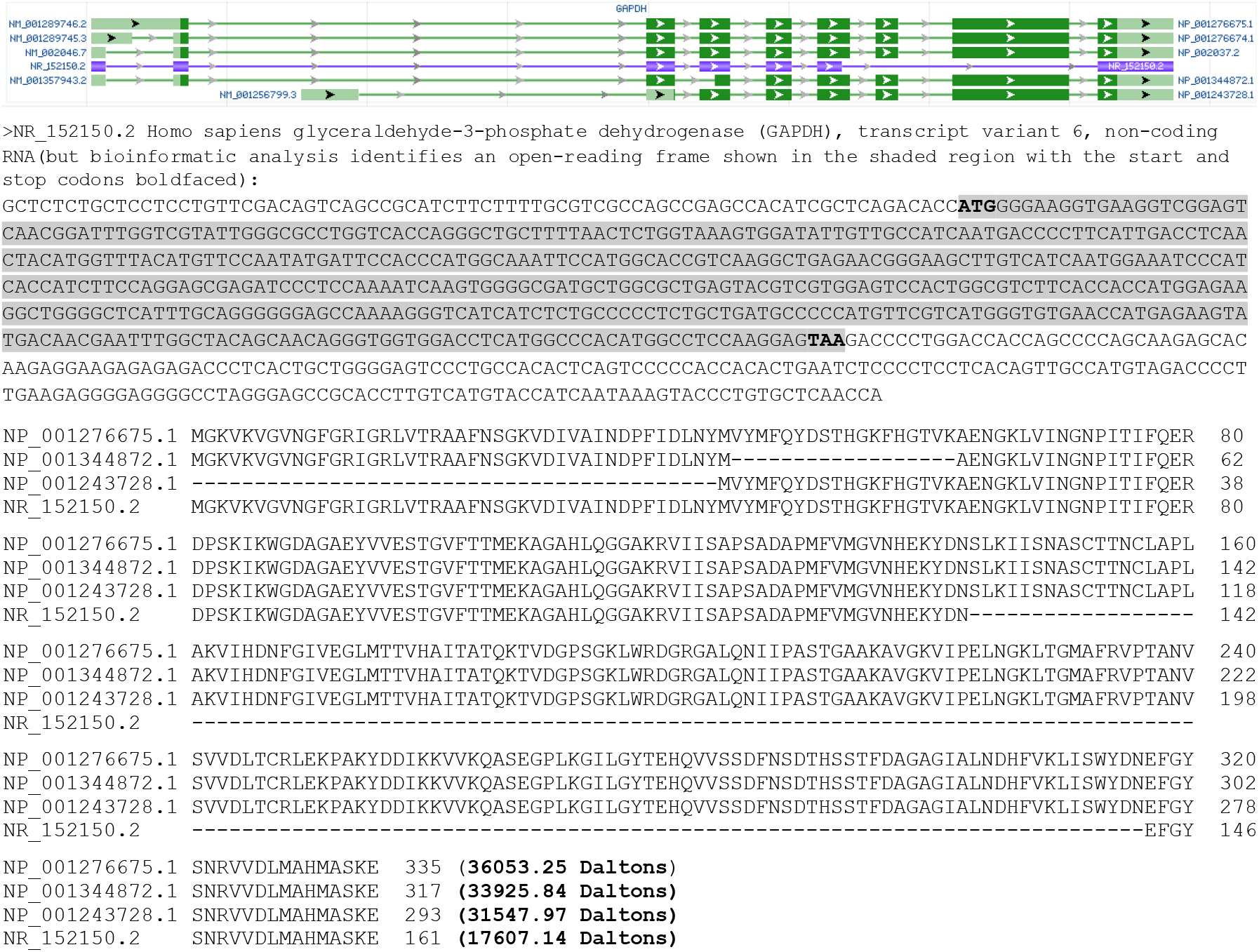
RNAs and proteins of the human GAPDH gene. An image copied from the NCBI database showing six RNA variants of GAPDH, of which NR_152150.2 is annotated as a non-coding RNA (Top panel) but, according to our analysis, it encodes an open reading frame that is shown as the shaded sequence with the start and stop codons boldfaced and encodes a GAPDH isoform of 161 AAs (Middle panel). Thus, the six RNAs encode a total of four protein isoforms, with their similarities and disparities as well as their molecular weights shown in the bottom panel.

NCBI lists only one ACTB RNA, which is an mRNA encoding a 375-AA protein (NP_001092.1). The “psms” datasheets (Supplementary tables 2 and 3) resulting from our LC-MS/MS analyses show that some identified ACTB peptides are shared by ACTC1 and ACTBL2, two other ACT family members, which is confirmed by the alignment of the protein sequences of ACTB, ACTC1, and ACTBL2 (Fig. 3). Moreover, the “psms” datasheets also show that some identified ACTB peptides are shared by the C-terminal region of the proteins of several POTE genes, i.e. POTEF, POTEE, POTEI, and POTEJ (Fig. 4, top panel). The POTE gene family has seven additional mRNA-encoding members, including POTEA, POTEB, POTEB2, POTEC, POTED, POTEG, and POTEM, besides the POTEKP that is a pseudogene coding for a non-coding RNA. The proteins of these seven POTE genes share only the N-terminal region with POTEF, POTEE, POTEJ, and POTEI, and thus do not have similarity to ACTB protein. Our conjecture is that the four longer POTE genes evolved by fusing one of the seven shorter genes at its 3’-end to the 5’-end of ACTB to form a fusion gene, which later evolved to the other three ACTB-containing POTE genes (Fig. 4, bottom panel). Interestingly, analysis of the evolutionary distances among the mRNAs of ACTB, ACTC1, ACTBL2, and the four POTE genes reveals that ACTB is evolutionarily closer to the four POTEs than to ACTC1 and ACTBL2 (Fig. 5). Therefore, it is likely that ACTB evolved to POTEF, POTEE, POTEJ, or POTEI later than to ACTC1 and even later to ACTBL2. In line with this conjecture, ACTB protein has a total of 39 AAs different from ACTC1 or ACTBL2 (Fig. 3) but only has 36 AAs different from one of the four TOPE proteins (Fig. 4).

**Fig. 3:**
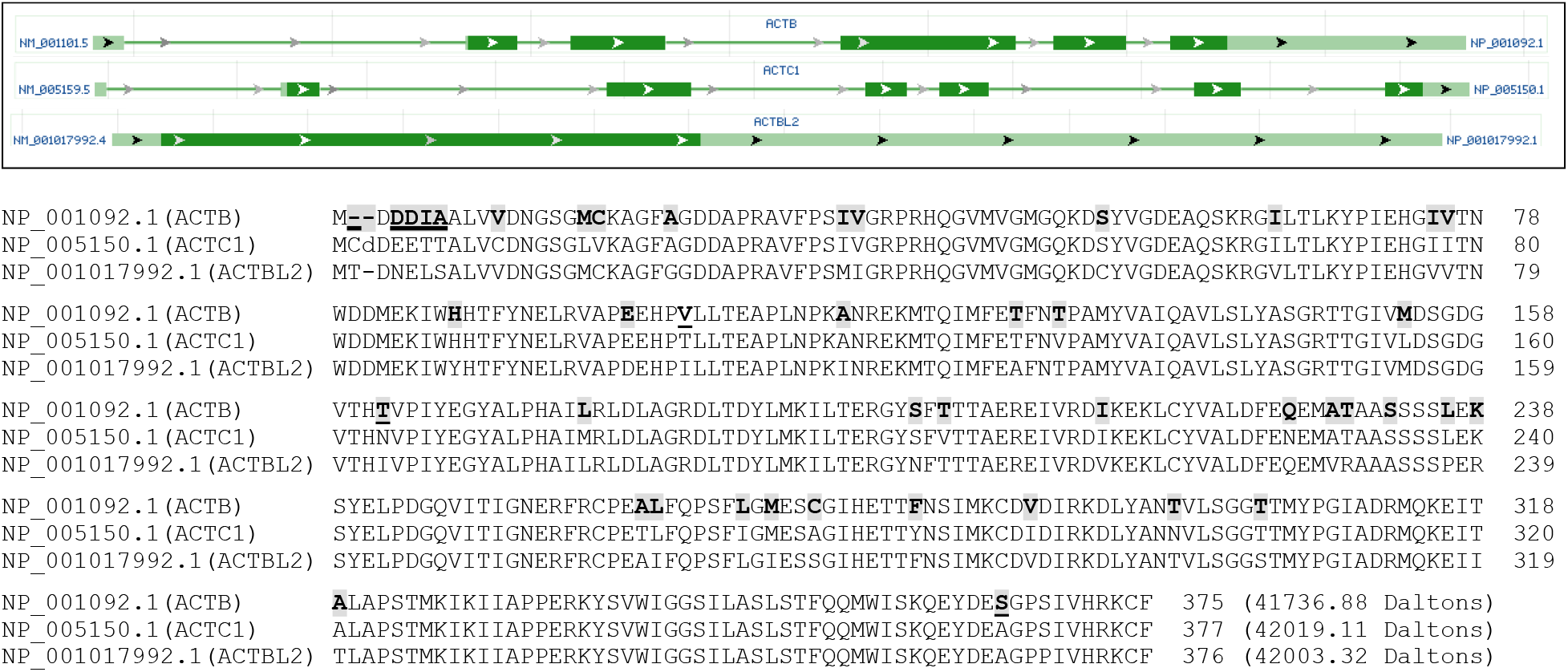
Similarities of ACTB to ACTC1 and ACTBL2. Images copied from the NCBI database show that ACTB, ACTC1, and ACTBL2 have only one RNA, with ACTBL2 being a one-exon gene (top panel). Alignment of the ACTB, ACTC1, and ACTBL2 proteins shows that they are highly similar in their AA-sequence (bottom panel). The AAs in ACTB that differ from either ACTC1 or ACTBL2 are shaded, whereas the AAs in ACTB that differ from both ACTC1 and ACTBL2 are shaded and underlined.

**Fig. 4:**
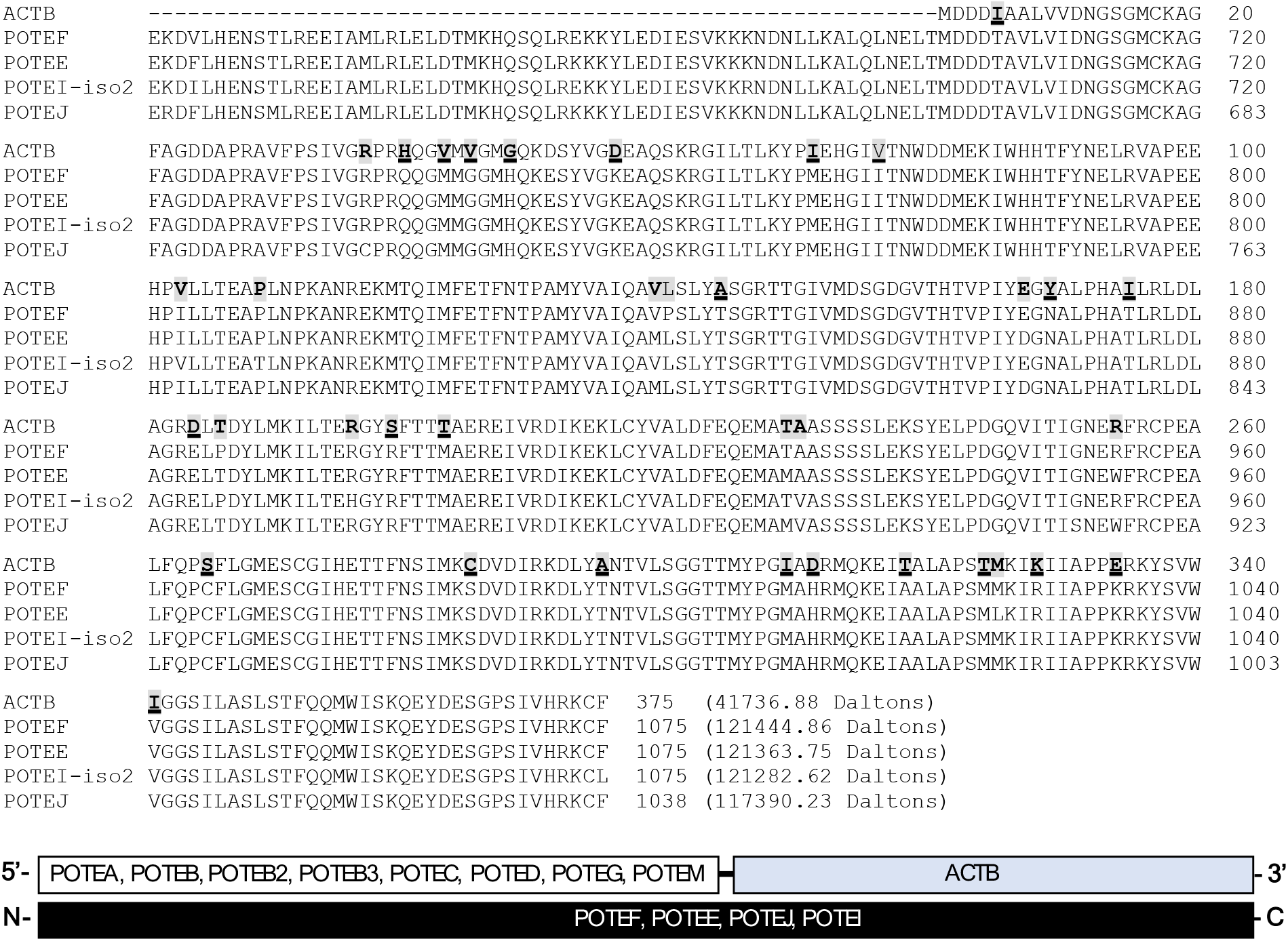
Alignment of the ACTB protein with the POTEF, POTEE, POTEJ proteins and the protein isoform 2 of POTEI shows that the ACTB protein is highly similar to the C-terminal region of these four POTE proteins (top panel). The AAs in ACTB that differ from only one, two, or three of the four POTE proteins are shaded, whereas the AAs in ACTB that differ from all of the four POTE proteins are shaded and underlined. We surmise that these four POTE genes might be formed as fusion genes between the 3’-end of the POTEA, POTEB, POTEB2, POTEB3, POTEC, POTED, POTEG, or POTEM gene and the 5’-end of the ACTB gene (bottom panel).

**Fig. 5:**
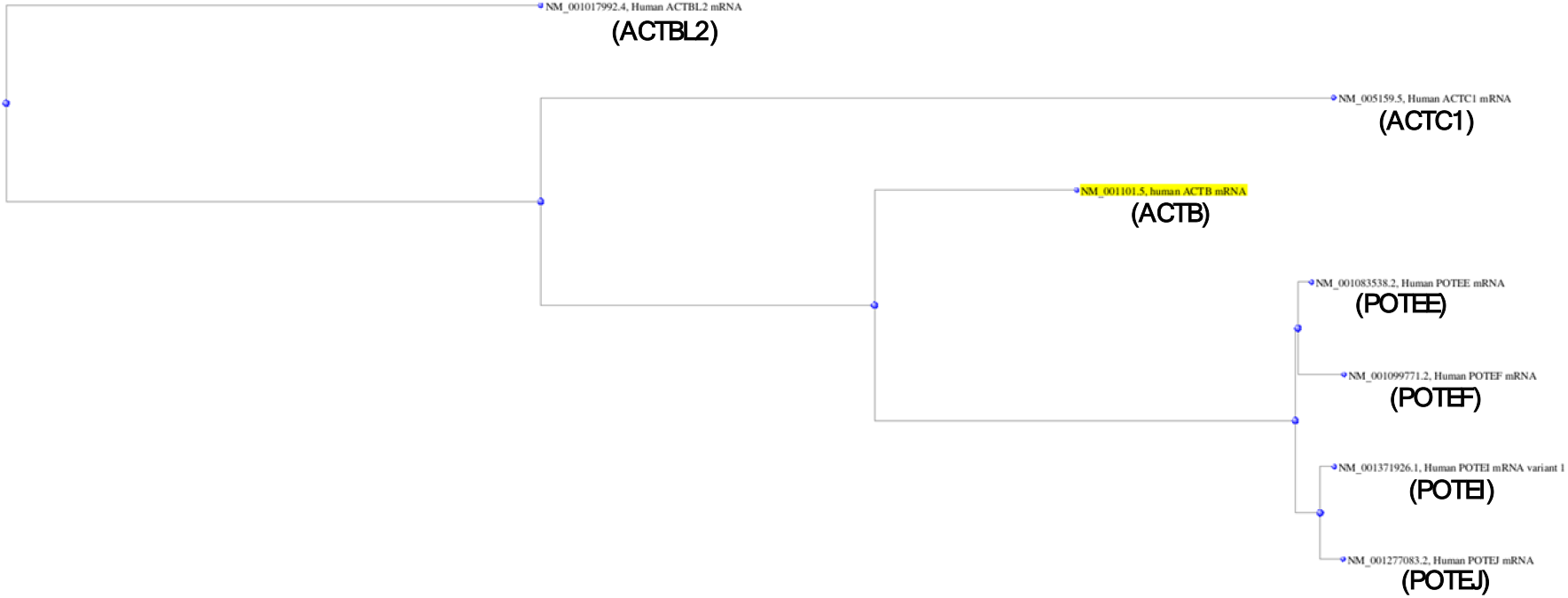
Distance tree resulting from analysis of the evolutionary distances among the mRNAs of ACTB (NM_001101.5), ACTC1 (NM_005159.5), ACTBL2 (NM_001017992.4), POTEF (NM_001099771.2), POTEE (NM_001083538.2), POTEI (NM_001277083.2), and POTEJ (NM_001371926.1).

### Detection of GAPDH and ACTB at the 72-, 55-, 48-, 40-, and 26-kD positions of SDS-PAGE

Although the full-length GAPDH is about 36 kD (Fig. 2), our LC-MS/MS analyses identified short peptides of GAPDH from both MB231 and MCF7 cell lines at the 72-, 55-, and 48-kD positions, from both MB231 and HEK293 cells at the 40-kD position, and from the HEK293 cells at the 26-kD position. All of the LC-MS/MS-identified peptides are unique to GAPDH, indicating that there is no protein that is significantly similar to GAPDH at the protein level and significantly expressed in these cell lines at any of these positions. We mapped all LC-MS/MS-identified peptides onto the full-length GAPDH protein and found that all four GAPDH isoforms should contain at least two unique peptides for LC-MS/MS identification (Fig. 6). We then calculated the coverage rate of each cell line at each gel position and found that all of the rates matched with the rates provided in the “proteingroups” datasheet (Table 1 and Supplementary table 1). Interestingly, the HEK293 cells at the lowest position, i.e. 26-kD, show the highest coverage rate, reaching 76.72% (Table 1 and Fig. 6). It is worth mentioning that, because the LC-MS/MS approach used short peptide(s) to predict the existence of a protein, the peptides detected in the same gel stripe may not necessarily belong to the same isoform, and it cannot be excluded that they belong to different known or unknown isoforms that have similar molecular weights and thus appear roughly at the same position.

**Fig 6:**
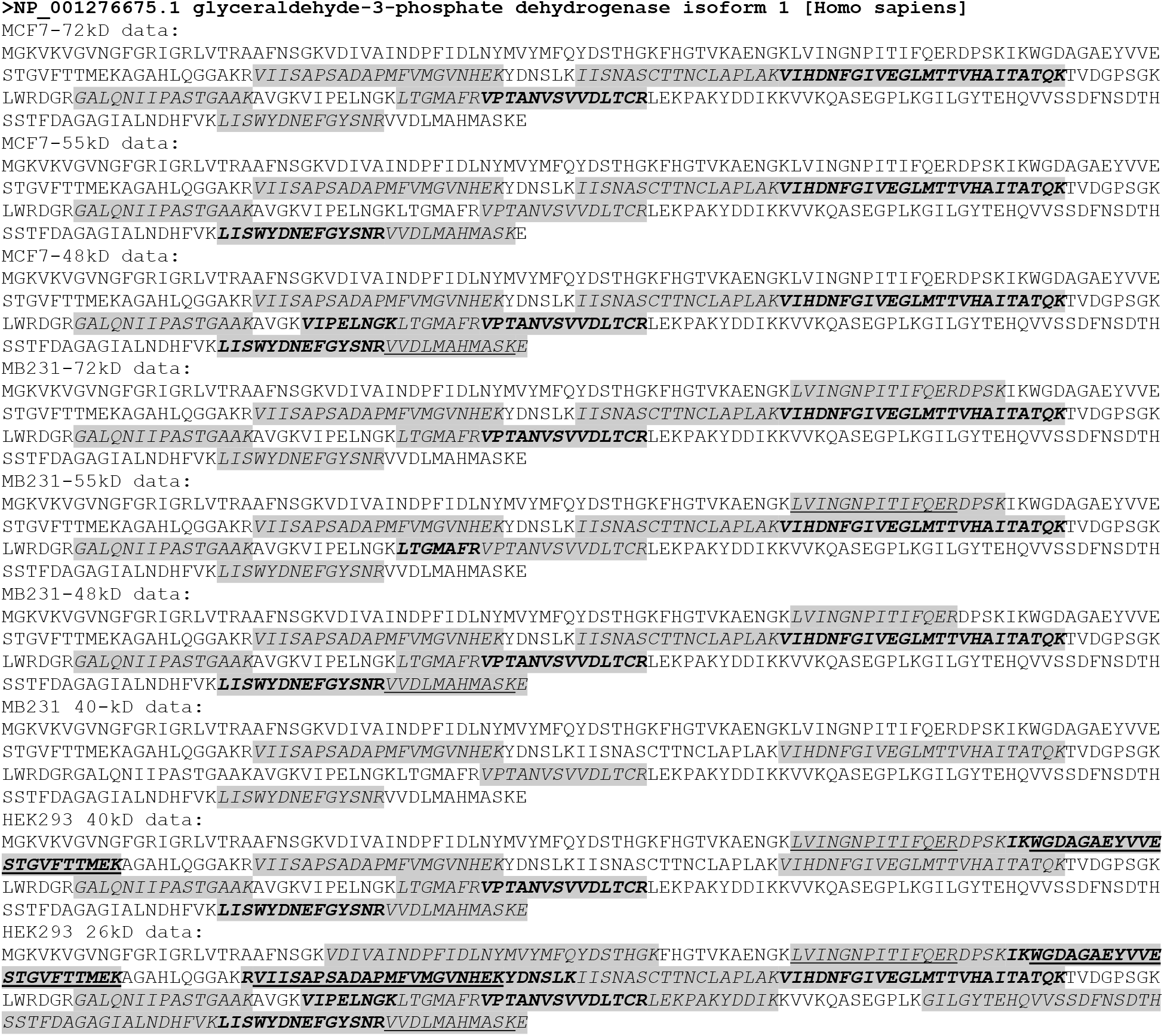
Mapping LC-MS/MS-identified peptides (shaded and italicized regions) onto the full-length GAPDH protein. Some long identified-sequences are actually formed by several consecutive identified peptides with boldfaced sequence(s) to segregate each peptide. Sometimes a peptide was identified as a slightly longer or shorter version of another peptide; in this case the shorter version is underlined. For instance, both “VVDLMAHMASKE” and “<u>VVDLMAHMASK</u>” are identified, with the underlined one lacking the “E”.

**Table 1:**
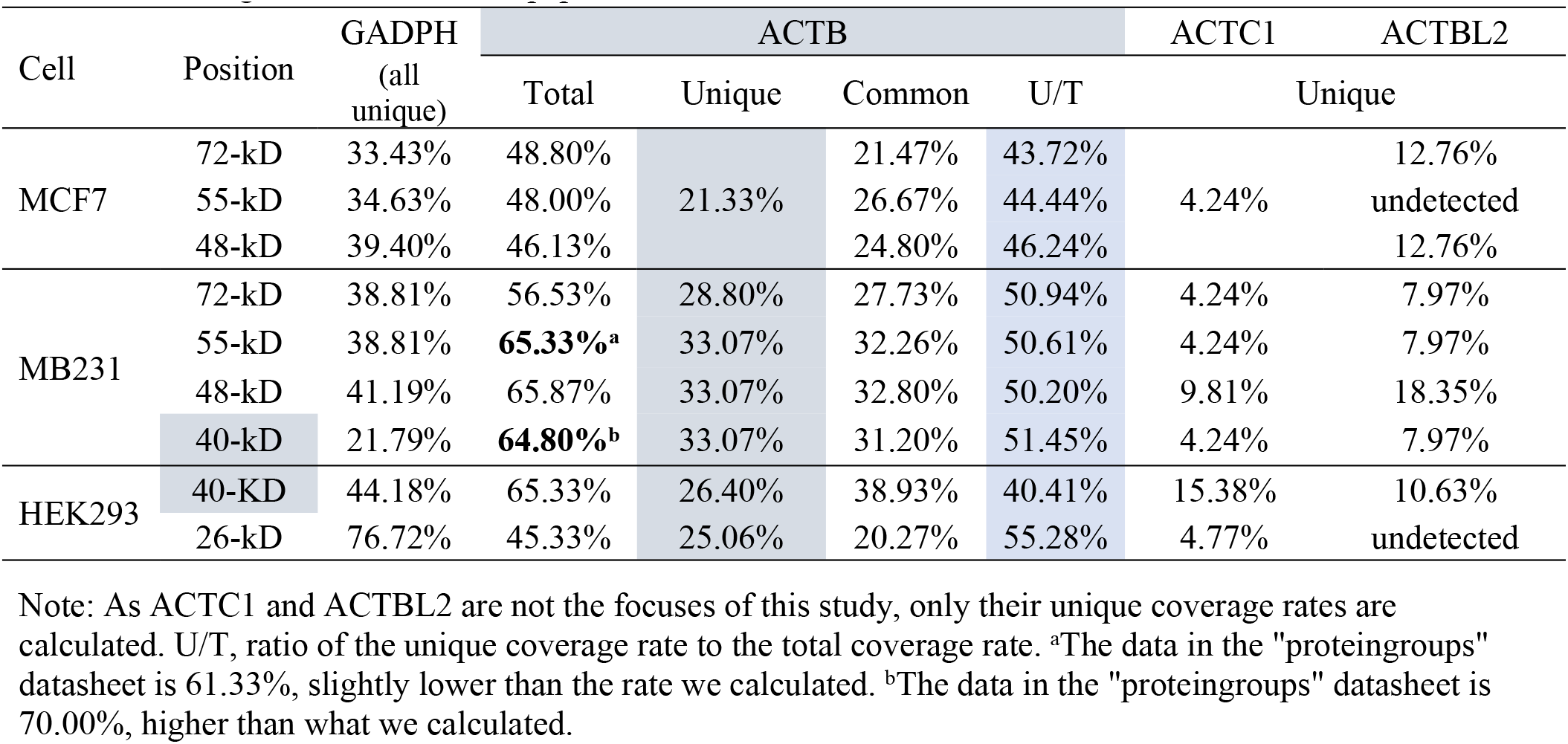
Coverage rates of identified peptides

Somewhat unlike the peptides of GAPDH, there are both unique and non-unique peptides of ACTB. The non-unique ones, referred to as “common” herein, are shared with the ACTC1, ACTBL2, POTEF, POTEE, or POTEJ protein, or with the protein isoform 2 of POTEI, and are the clues leading us to the discovery of the similarity of ACTB to ACTC1 and ACTBL2 as well as the discovery of the four ACTB-containing POTE genes. We mapped all identified peptides onto the full-length ACTB protein and calculated the coverage rates by the unique peptides and the total coverage rate by both common and unique peptides (Fig. 7). Most of the total coverage rates matched the rates given in the “proteingroups” datasheet (Table 1 and Supplementary table 1). However, for unknown reasons two of the coverage rates in our calculations differ slightly from the figures given in the “proteingroups” datasheet (65.33% *vs* 61.33% and 64.80% vs 70.00%; Table 1). Notwithstanding, the total coverage rates of the identified peptides are high for different cell lines at different SDS-PAGE positions; the unique coverage rates are also high, varying between 21.33% and 33.07%, and contribute to more than 40% of the total coverage rates (Table 1).

**Fig 7:**
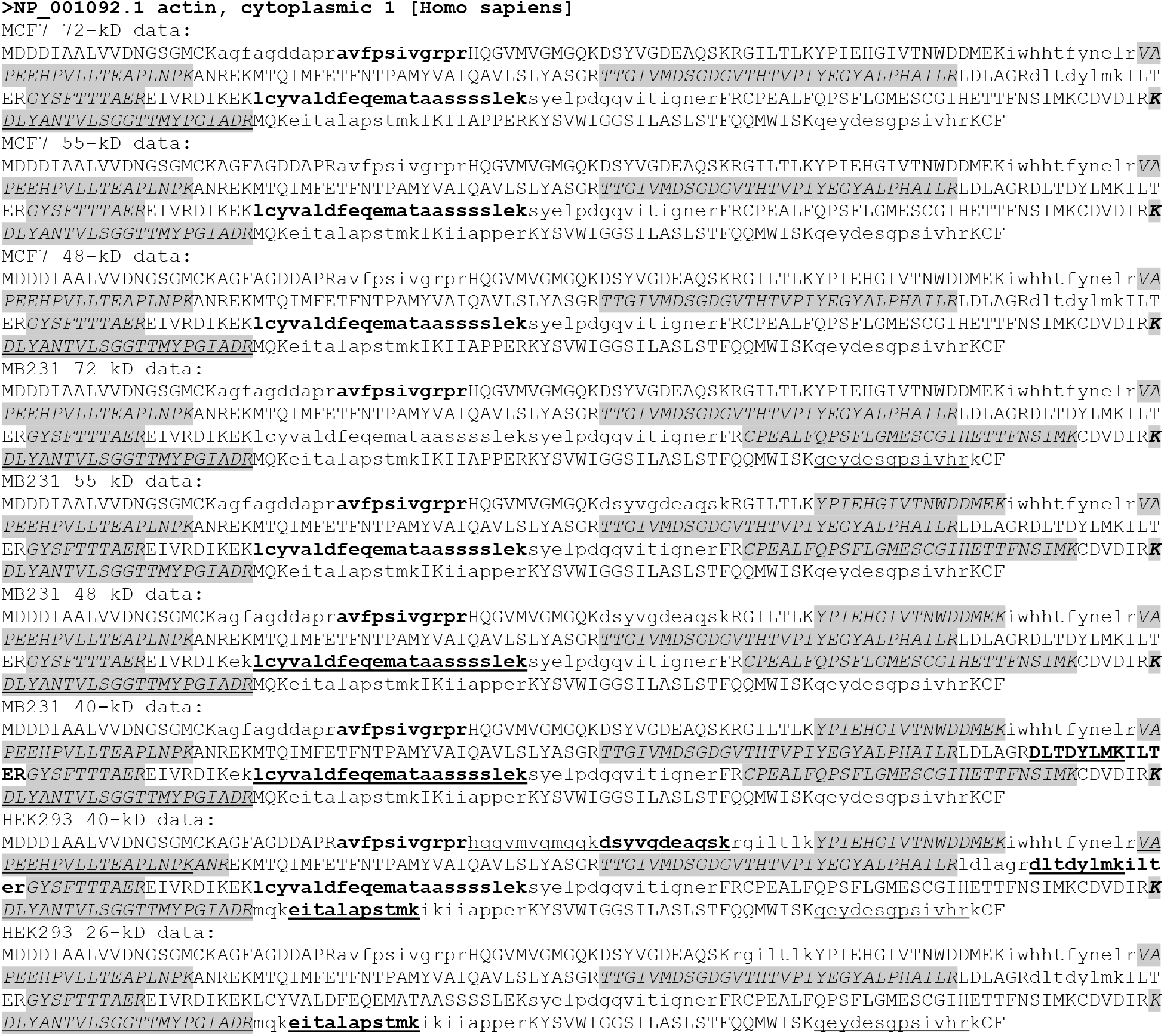
Mapping LC-MS/MS-identified peptides onto the full-length ACTB protein, with the shaded and italicized regions being the unique peptides and the lowercase regions being the common peptides. Some long identified-sequences are actually formed by several consecutive identified-peptides with boldfaced sequence(s) to segregate each peptide. Sometimes a peptide was identified as a slightly longer or shorter version of another peptide; in this case the shorter version is underlined. For instance, both “QEYDESGPSIVHRK” and “<u>QEYDESGPSIVHR</u>” are identified, with the underlined one lacking the “K”.

Some peptides, and some AAs in a peptide, are identified in some cell lines at some SDS-PAGE positions but not in or at some others. We counted those AAs that have been identified in at least one cell line at one position to obtain the theoretical maximal-identified AAs, which is 252 AAs for ACTB. Because the ACTB protein has 375 AAs, its theoretical maximal-total-coverage rate is 252/375, i.e. 67.20%. None of the cell lines at any position shows a total coverage rate as high as this theoretical maximum, but many of the rates we calculated are close to it (Table 1). In a similar way we obtained the theoretical maximal-unique-coverage rate for ACTB, which is 33.07% and has actually been obtained in the MB231 and HEK293 cells for most positions but not in the MCF7 cells at any position (Table 1), likely due to some technical reasons.

### Some possible translational and post-translational mechanisms for generation of isoforms

As we frequently discussed before [2,26–29], utilization of a downstream start codon in a mRNA for translation to generate a protein isoform with a shorter N-terminus is very common, with the generation of some smaller isoforms of c-Myc, P53, and RB as good examples [30–32]. Theoretically, this mechanism may also be used in translation of GAPDH, ACTB, POTEE, POTEF, POTEI and POTEF to generate isoforms with a shorter N-terminus, since all these genes have many ATGs in the same ORF with the canonical one, with POTEF shown in Figure 8 (top panel) as an example. Other inframe start codons besides ATG, such as CTG that is used for translation of a c-Myc or PTEN isoform [30,33], may also exist but are not analyzed as they are less used. Moreover, physiological single nucleotide polymorphisms and pathological genetic alterations such as single nucleotide mutations may alter the canonical start codon; in this case, translation may also be incepted with a downstream start codon as described above. If there is an upstream ORF, its translation may be extended to the annotated ORF at its downstream, thus engendering a longer N-terminus (Fig. 8, bottom panel). On the other hand, if such polymorphism or mutation occurs at the stop codon, translation may be extended to a downstream one, resulting in an isoform with C-terminal extension (Fig. 8, top panel).

**Fig. 8:**
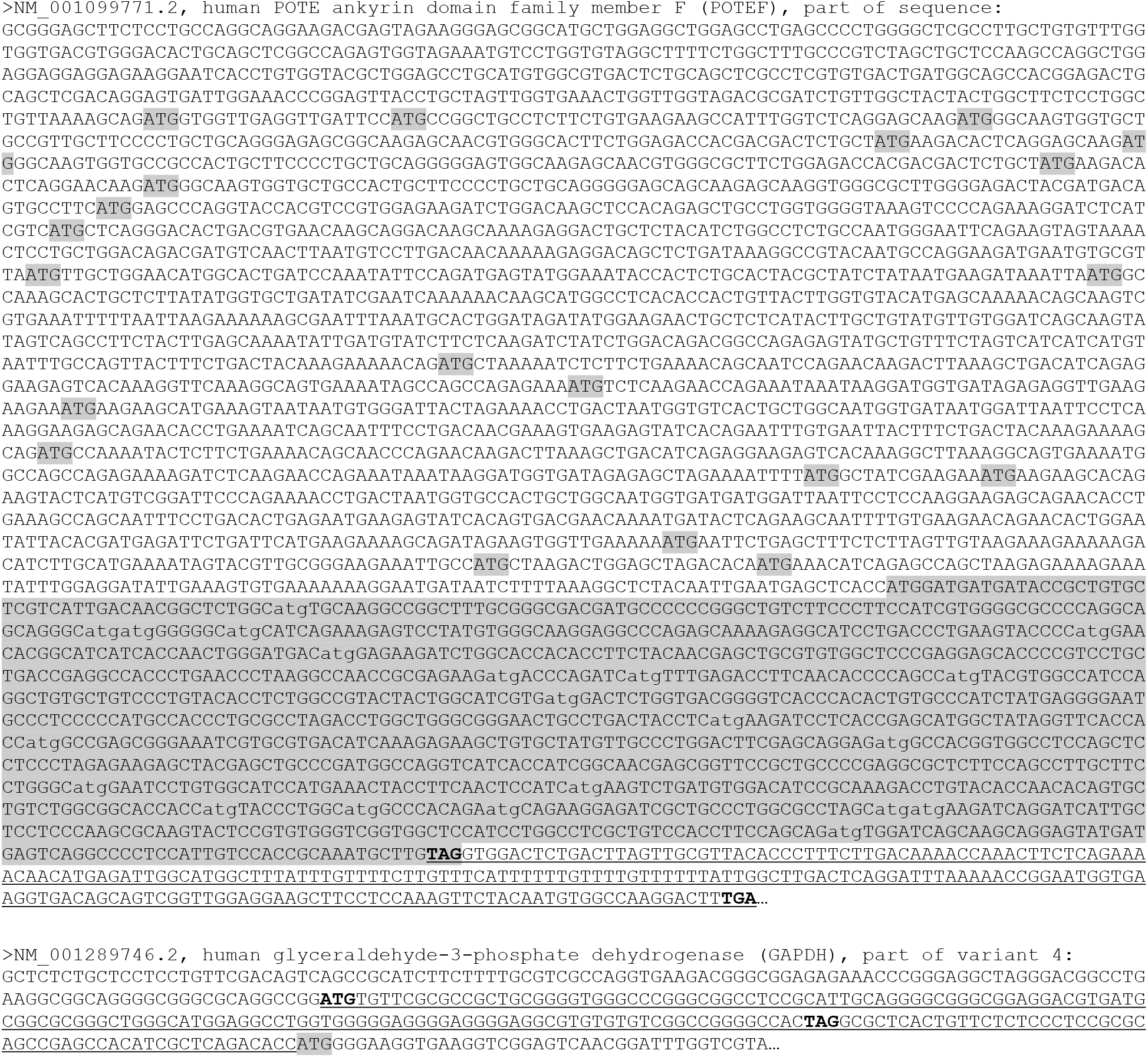
Depiction with examples of some mechanisms for N- or C-terminal extension of a protein caused by a mutation and for N-terminal truncation caused by use of a downstream ATG. **Top-panel**: Part of POTEF mRNA sequence, with all in-frame ATG start codons and the ACTB-homologous region shaded. If translation starts with any of the downstream ATGs, a smaller POTEF isoform will be generated that may be mistaken as a larger ACTB isoform on a WB membrane with an ACTB antibody that cross-reacts with POTEF. On the other hand, if a mutation occurs in the TAG stop codon (boldfaced), translation will be extended to a downstream TGA stop codon (boldfaced), producing a POTEF isoform with 73 more AAs at the C-terminus encoded by the underlined sequence, which may also be mistaken as an ACTB isoform in WB by a cross-reactive antibody. **Bottom-panel**: Part of the 5’-sequence of a GAPDH mRNA showing an upstream ORF that is in-frame with the annotated ORF. If a mutation occurs in the TAG stop codon (boldfaced) of this upstream ORF, its translation initiated from the upstream ATG (boldfaced) will be extended to the annotated start codon (shaded), producing an isoform with 64 more AAs at the N-terminus encoded by the underlined sequence.

After translation, proteins are often subjected to many different types of chemical modification that can affect their migration in SDS-PAGE. We therefore calculated the changes in molecular mass that may be caused by some common chemical modifications (Table 2). For instance, one cholesterolation, glycosylation, GPI (glycosylphosphatidylinositol) anchor, ubiquitination, and SUMOylation can, theoretically, increase about 0.4 kD, 0.45-3.3 kD, 2-3 kD, 8.6 kD, and 12 kD, respectively, while some other types of chemical modification change the molecular mass only slightly (Table 2). Many of these chemical modifications, such as phosphorylation, can simultaneously occur to many AAs of a protein, collectively making a huge impact on migration of the protein. It is worth mentioning that some chemical modifications, such as phosphorylation, not only can change the molecular mass but also may alter the electronic charge of the protein, and thus may accelerate or decelerate protein migration in SDS-PAGE. Moreover, polyubiquitin, and poly-SUMO, polyglycylation, polyglutamylation, polyamination, etc., can occur as a chain, most of which have been well studied for tubulin as an example [33–35]. Any of these chains can greatly slow down protein migration.

**Table 2:**
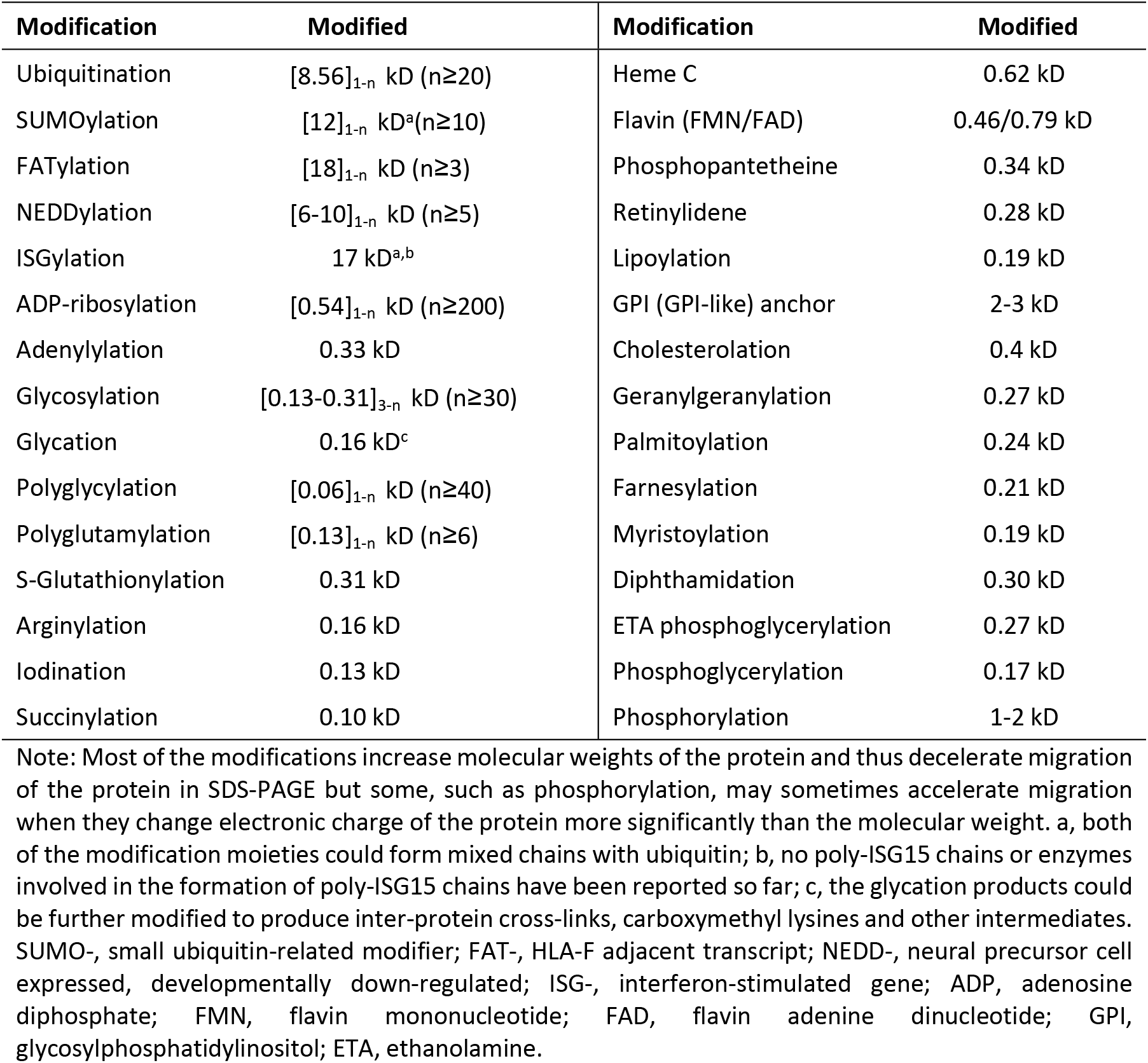
Some post-translational chemical modifications of proteins that affect protein migration in SDS-PAGE:

## Discussion

In our previous studies, we observed unexpectedly that most proteins from several human cell lines detected at the 72-, 55-, 48-, 40- and 26-kD positions of SDS-PAGE had a theoretical molecular mass that is either too large or too small for them to appear at the detected position [22– 24]. Even more unexpected is that many proteins were detected simultaneously at two or more positions [22–24], just as what we are reporting herein for GAPDH and ACTB. Considering that most proteins could indeed be detected with WB at the expected position of SDS-PAGE, as reported in numerous publications by numerous researchers, including us, we then concluded that most genes might express at least one additional protein isoform besides the WT or the canonical form that was also detected by us [22,23]. This conclusion is congruent with today’s knowledge that most human genes can engender multiple protein isoforms [2,36], although a general estimation has not yet been available on the total number of the genes with protein multiplicity in the human genome. However, this conclusion did not consider another possibility: since LC-MS/MS uses short peptide(s) to predict the existence of a whole protein, some detected peptides may not be derived from the authentic genes but, instead, are derived from other currently-unknown genes that contain the elements of the authentic genes. By analyzing our LC-MS/MS-identified peptide sequences, we inadvertently found in the current study the existence that POTEE, POTEF, POTEI, and POTEJ genes have a region highly similar to ACTB, which strengthens this possibility. There are indeed a huge number of unknown or unannotated genes in the human genome as shown in NCBI database, and genome-wide analyses of many gene-knockout mice also suggests the existence of a huge number of unannotated genes in the mouse genome [37]. In line with this possibility, there are a large number of unmatchable MS data in our previous studies, some of which may be derived from unannotated peptides [22,23]. Moreover, under pathological situations, new fusion genes containing ACTB or GAPDH elements may be formed, with the ACTB-FOSB and ACTB-GLI1 fusion genes in some neoplasms as instructive examples [9–13]. More intricately, the concept of “unknown gene” may be defined at the levels of RNA and even cistron as well, and not necessarily at the chromosomal DNA level. This is because an unknown protein can be generated from an additional cistron of a mRNA encoding the canonical protein, since it has been estimated that the number of polycistronic mRNAs engendered from the human genome is colossal [38]. Many non-coding RNAs contain one or more cistrons as well [39–41], with GAPDH’s non-coding RNA NR_152150.2 shown herein as a paradigm. It is also worth mentioning that NCBI database updates its data almost on a daily basis, often by adding more new RNA variants or protein isoforms to many genes’ datasets. For instance, there was only one GAPDH RNA listed in NCBI a few years ago when we studied GAPDH pseudogenes [1], but now there are six.

The GAPDH and ACTB detected at the 40-kD position may be the WT form of 36 kD and 41.7 kD, respectively, as protein migration in an SDS-PAGE gel can be affected by various factors and the pre-stained protein markers are not very accurate. The ACTB, and even the GAPDH, detected at the 48-kD position may still be the WT form as well, if they have been subjected simultaneously to multiple post-translational modifications. A combination of multiple chemical modifications listed in Table 2 with the aforementioned variations in protein migration and in the pre-stained markers can significantly affect the appearance of a protein on the gel and can even, theoretically, shift GAPDH and ACTB to the 55-kD and 72-kD positions. Even the same type of chemical modification that occurs in a chain, such as a chain of polyubiquitin, poly-SUMO, polyglycylation, polyglutamylation, or polyamination, can greatly decelerate protein migration in SDS-PAGE as well. However, there are other possibilities besides chemical modifications: first, the GAPDH and ACTB detected at the 48-, 55-, and 72-kD positions are unknown isoforms that are larger than the canonical form. Second, the peptides detected at these positions are derived from unannotated genes evolving from GAPDH or ACTB as already described above. The possible existence of multiple protein isoforms of GAPDH dovetails with its functional versatility [9–13].

A simple assumption for the appearance of GAPDH and ACTB at the 26-kD position is that some of their molecules have degraded randomly. Both GAPDH and ACTB are abundantly expressed; accordingly, random degradation may occur more often. LC-MS/MS-detected peptides are distributed throughout the whole protein sequence of GAPDH or ACTB with a very high coverage rate, which is also in congruence with random degradation. However, there still are other possibilities: first, GAPDH and ACTB may express one or more currently unknown isoforms that are smaller than the canonical form, such as via utilization of a downstream translation start codon for translation of an isoform with a shorter N-terminus, as depicted in Figure 8. Actually, if this scenario occurs to one of the four TOPE proteins as well, it will produce a smaller isoform with a molecular weight varying from several kDs to 120 kDs (the molecular weight of the canonical POTE), which may be mistaken as an ACTB protein in WB using a POTE antibody that cross-reacts with ACTB. It needs to be mentioned that detection of proteins at a position much lower than the proteins’ theoretical molecular mass, like the detection of GAPDH or ACTB at 26-kD, has been discerned in our previous studies for a large number of genes’ protein products [22,23]. It is likely that different mechanisms may be used for generation of different protein isoforms of different genes.

Besides the above-described scenarios that may occur physiologically, in many pathological situations, including in immortalized cell lines, mutations may occur leading to generation of larger or smaller protein isoforms of a gene via different mechanisms. For example, GAPDH has several upstream start codons and stop codons in its RNA variants that are in-frame with the annotated ORF (Fig. 8). If a single nucleotide mutation occurs to an upstream stop codon, translation initiated from an upstream ATG may be extended to the annotated ORF, resulting in an N-terminal-extended GAPDH isoform (Fig. 8). Similarly, if a mutation occurs to the annotated stop codon, translation will be extended to a downstream stop codon, resulting in a C-terminal extended isoform (Fig. 8).

For both ACTB and GAPDH, some of the peptides were detected in some cell lines at some positions but not in or at some others, as shown in Figures 6 and 7. The reasons are multiple and could be technical or biological (physiological or pathological). For instance, a higher abundance of a protein will provide a higher chance for its peptides to be detected in a LC-MS/MS procedure. Ionic strength of a peptide is the basis for its detection by MS, but it often differs greatly among different peptides, creating differences among different peptides in the chance of being detected. However, among all possible explanations exists a simple one: the absence of a peptide that has been detected in other cell type(s) at other position(s) connotes that the particular isoform lacks this peptide sequence. Therefore, identification of the missing peptide(s) by mapping of the detected peptides onto the gene’s protein sequence, such as what we did for the GAPDH or ACTB protein shown in Figures 6 and 7, may provide us with clues for further identification of unknown isoforms of the gene being studied that have a specific region deleted.

Because most human genes can produce multiple protein isoforms, it should be very often that WB detects not only the expected band but also additional band(s) on the membrane. In reality, both are common as WB indeed detects additional band(s) at “wrong” position(s) on the membrane and as WB detects only the expected band. In the former situation, it is a common but hardly mentioned practice for the researchers to assume, although without supporting evidence, that those bands at “wrong” positions are non-specific proteins and should be cut off, presenting in the publications only the band of interest. It is very often that antibody supplier companies are blamed for selling “lousy, not specific enough” antibodies. To avoid being blamed, the suppliers try their best to select and supply those antibodies that recognize only a single band at the expected molecular weight on WB membranes, which usually is the WT or the canonical protein form. This is technically feasible since different isoforms having the same antigen-sequence may manifest different conformations inside the antibody-producing animal, making B lymphocytes produce some antibodies that recognize only one isoform but not the others. As we have discussed before [36,42], this compromise between the researchers and the antibody suppliers actually leads to a biased, somewhat misleading, result. Many published WB results that have only a single band detected may be due to this compromise, although there certainly are many cases in which the gene of interest does indeed produce only a single protein form in the given cell types at the given situation. Although primary antibodies that recognize only a single isoform are useful, those that recognize multiple isoforms and thus seemingly are not specific enough may provide us with a more global picture of the protein products of the gene studied. For this reason, we suggest anew to peers to retain and present all bands appearing on the WB membrane unless there is tangible evidence proving that the additional band(s) are non-specific.

We [1] and many RNA pundits have proffered that ACTB and GAPDH are not suitable for serving as the reference genes for determination of the RNA level [3–8,43–50], and have pointed out that it is a fallacy to have a universal reference gene for RT-PCR and that GAPDH or ACTB is certainly not the one [6–8,51–55]. Unfortunately, the vast majority of biomedical researchers keep ignoring this advice and continue using either GAPDH or ACTB as the only reference. We herein further proffer that GAPDH and ACTB should not be used either as the references for determination of the protein level in such techniques as WB. The reasons for the unsuitability of GAPDH are threefold: first, GAPDH has at least three additional protein isoforms, including the one found herein encoded by an RNA variant annotated by NCBI as a non-coding RNA. Should a researcher intend to use GAPDH, it needs to be determined first whether any of the three non-canonical isoforms is expressed in the target cells in the situation of interest. Probably, it is easier to find another reference gene instead of performing such tedious prerequisite work. Second, our detection of GAPDH peptides at multiple positions of SDS-PAGE suggests that, either there are unknown GAPDH isoforms or unknown GAPDH-containing proteins from unknown genes, or some of the known GAPDH isoforms shown in Figure 2 have experienced complicated chemical modifications, likely some of those listed in Table 2 but uncharacterized for GAPDH. Although there are commercial antibodies that recognize only the canonical form, one may not be able to claim that it represents the real profile of GAPDH protein expression. Third, GAPDH has too many different functions. The great similarity of ACTB to ACTC1 and ACTBL2 is an evident reason for its unsuitability as a protein reference, besides the first two of the aforementioned reasons for GAPDH.

In summary, we found multiple peptides of ACTB and GAPDH at multiple SDS-PAGE positions, which raises a few questions, such as whether these two genes express some unknown protein isoforms. GAPDH has four known protein isoforms, including one encoded by an RNA variant annotated by NCBI as a non-coding RNA, whereas ACTB is highly similar in AA sequence to ACTC1, ACTBL2, and proteins of four POTE family members. These data, together with the known fact that GAPDH has versatile functions, lead us to a somewhat provocative conclusion that ACTB and GAPDH are not suitable for serving as the reference genes for determination of the protein level, a leading role these two genes have been playing for decades in the biomedical research.

## Authors’ contributions

YH drafted the manuscript. JZ and SL performed the LC-MS/MS and analyzed the data. JQ analyzed the data and prepared the figures and tables. YZ participated in the discussion and manuscript revision. LZ performed the English editing of the manuscript and participated in the discussion. HH, JZ and DZL conceptualized the study. DZL performed the SDS-PAGE and gel stripe excision and finalized the manuscript.

## Acknowledgements

We would like to thank Dr. Fred Bogott at Austin Medical Center, Mayo Clinic in Austin, Minnesota, USA, for his excellent English editing of this manuscript.

## Ethics approval and consent to participate

not applicable.

## Consent for publication

All authors agree on the publication.

